# Precision Through Frequency-Aware Prioritization: Introducing the Bouziane Similarity Index (BSI) for Rare Variants

**DOI:** 10.1101/2025.04.28.651048

**Authors:** Ismail Bouziane, Khelifa Louafi, Okba Louafi, Meriem Allouche, Omar Sekiou, Mahieddine Boumendjel

## Abstract

Current genetic similarity metrics, including Identity-by-State (IBS) and Kinship coefficients, treat all alleles equally, ignoring allele frequency differences and partial genetic sharing (Purcell et al., 2007; Speed & Balding, 2015). Despite the recognized biological importance of rare variants (Lee et al., 2012), no widely adopted similarity metric prioritizes rare variant sharing in relationship inference.

We introduce two novel metrics: the Bouziane Similarity Index (BSI), a frequency-weighted score emphasizing rare variant matches, and BSI-VAR, a variant that integrates allele dosage and inverse variance weighting to capture partial allele sharing and amplify rare variant signals. To evaluate these metrics, we first performed unsupervised principal component analysis (PCA) on simulated genotype similarity matrices, showing that BSI and BSI-VAR provide clearer genetic structure compared to IBS. We then conducted controlled simulations involving Parent-Child, Sibling, and Random genetic pairs to quantify discriminative performance.

Through all evaluations, BSI and especially BSI-VAR consistently outperformed IBS in distinguishing genetic relationships. These findings establish BSI and BSI-VAR as robust, interpretable, and rare-variant-aware alternatives for modern genetic similarity analysis, with potential applications in population genetics, ancestry inference, and precision medicine (Ashley, 2016; Manolio et al., 2009).

## Introduction

Accurately quantifying genetic similarity between individuals is essential across a range of genomic applications, including ancestry inference, population genetics, relationship mapping, and forensic analysis (Purcell et al., 2007; Speed & Balding, 2015). Traditional similarity metrics, such as Identity-by-State (IBS) and Kinship coefficients, have long served as foundational tools. However, these approaches have intrinsic limitations that reduce their resolution and sensitivity, particularly when applied to structured populations or in the presence of rare variants (Speed & Balding, 2015). IBS metrics simply count allele matches between individuals without accounting for allele frequency (Purcell et al., 2007). Consequently, common variants—which are frequently shared among unrelated individuals—contribute equally to similarity scores as rare, highly informative variants (Weir et al., 2006). This inflates genetic similarity estimates and diminishes the ability to distinguish between truly related and unrelated individuals. Similarly, Kinship coefficients estimate the probability of sharing alleles identical by descent (IBD) but rely on assumptions of random mating and Hardy-Weinberg equilibrium, which are often violated in real-world datasets (Speed & Balding, 2015; Weir et al., 2006). Although the biological importance of rare variants is well established, particularly in disease association studies and evolutionary biology (Lee et al., 2012), no widely adopted similarity metric explicitly prioritizes rare variant sharing in measuring genetic relatedness (Purcell et al., 2007; Speed & Balding, 2015). Furthermore, precision medicine initiatives increasingly depend on the detection of rare variants to inform diagnosis, prognosis, and therapeutic strategies (Ashley, 2016; Manolio et al., 2009), underscoring the need for frequency-aware genetic similarity measures.

To address these limitations, we introduce two novel metrics: the Bouziane Similarity Index (BSI) and its advanced variant BSI-VAR. BSI incorporates frequency-aware weighting, emphasizing matches at rare variants through inverse allele frequency scoring. BSI-VAR extends this framework by integrating allele dosage differences and variance-based weighting, enabling partial match scoring and capturing subtler genetic relationships.

We evaluated these metrics through unsupervised principal component analysis (PCA) of simulated genotype similarity matrices, demonstrating that BSI and BSI-VAR provide clearer global genetic structure compared to IBS. Controlled simulations involving Parent-Child, Sibling, and Random genetic pairs further confirmed that BSI and especially BSI-VAR consistently outperform IBS in discriminating genetic relationships. These results establish BSI and BSI-VAR as robust, interpretable, and rare-variant-aware alternatives for modern genetic similarity analysis.

## Methods

### 1. Metric Definitions

#### 1.1 Bouziane Similarity Index (BSI)

The Bouziane Similarity Index (BSI) quantifies genetic similarity between two individuals by prioritizing rare variant matches using frequency-aware weighting.

For individuals and across SNPs:

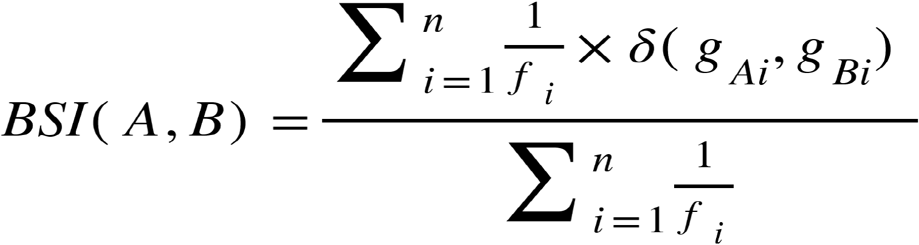

where:

- *i* indexes SNPs (*i* = 1, 2,…, *n*),
- *n* is the total number of SNPs,
- *f*_*i*_ is the minor allele frequency (MAF) of SNP*i*,
- *g*_*Ai*_ and *g*_*Bi*_ are genotypes of individuals *A* and *B* at SNP*i*,
- δ(*g*_*Ai*_ *g*_*Bi*_) = 1 if *g*_*Ai*_ = *g*_*Bi*_, and 0 otherwise.

The design of *BSI* ensures that matches at rare variants contribute more heavily to the overall similarity score than matches at common variants.

### 1.2 Bouziane Similarity Index with Variance Adjustment (BSI-VAR)

#### 1.2.1 Definition and Objective

The Bouziane Similarity Index with Variance Adjustment (BSI-VAR) is designed to quantify genetic similarity between two individuals by simultaneously:

- Prioritizing rare variant matches,
- Proportionally penalizing partial mismatches,
- Integrating dosage sensitivity through allele counts.

Unlike traditional methods such as Identity-by-State (IBS), BSI-VAR provides a frequency-aware and variance-aware approach, ensuring greater resolution and biological interpretability.

#### 1.2.2 Mathematical Derivation

Given two individuals *i* and *j*, and a set of *K* SNPs indexed by *k*, we define:

- *x*_*ik*_ = dosage of minor allele (0, 1, 2) for individual *i* at SNP *k*,
- *x*_*jk*_ = dosage of minor allele for individual *j* at SNP *k*,
- *p*_*k*_ = minor allele frequency (MAF) of SNP *k* in the population.

The local similarity at SNP *k* is defined as:

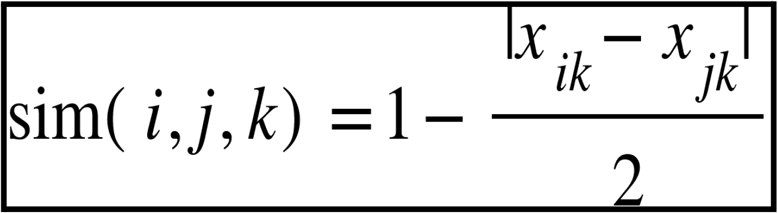

Each SNP is assigned a ***weight*** based on its variance under binomial expectation:

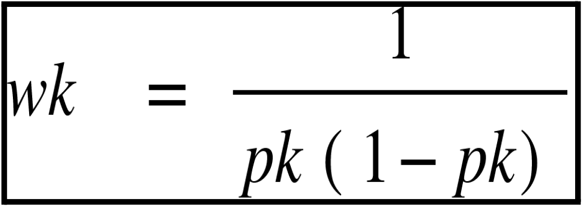

The **raw BSI-VAR score** between individuals *i* and *j* is the weighted sum:

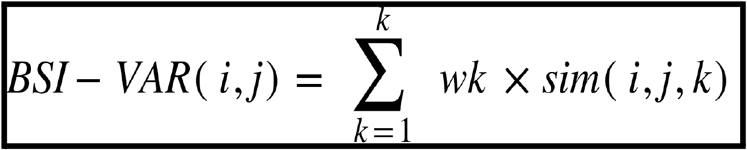

To normalize the score between 0 and 1:

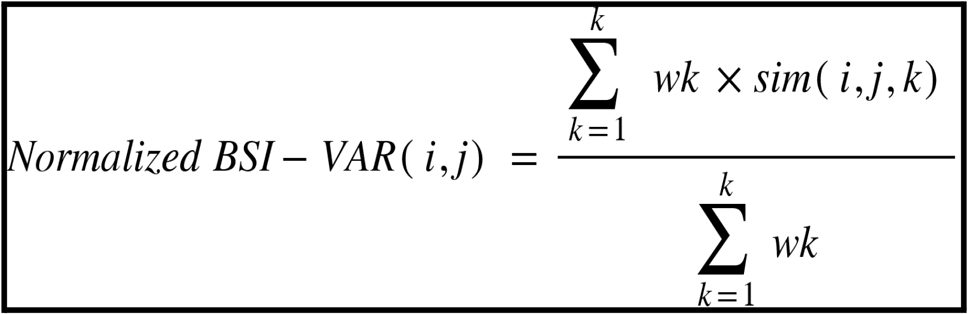

***Note***: *The normalized BSI-VAR is not simply sim(i, j, k*), *it represents the weighted average of local similarities sim(i, j, k*) *across all SNPs, scaled by the total weight. If only a single SNP is considered, Normalized BSI-VAR reduces to sim(i, j, k*) *at that locus*.

### 2.1 Tools and Software

All simulations, similarity score calculations, and visualizations were performed using Python 3.10.

The following Python libraries were utilized:

NumPy (v1.24.0) for numerical operations and array manipulation,Pandas (v1.5.3) for genotype data structuring,Matplotlib (v3.6.2) for data visualization and plotting,Scikit-learn (v1.2.1) for Principal Component Analysis (PCA), SciPy (v1.10.0) for statistical testing (Mann-Whitney U tests),tqdm (v4.64.1) for simulation progress tracking.

Simulations were run locally on a standard personal computer (Intel Core i5 processor, 8 GB RAM) without the need for specialized hardware.

### 2.2 Reproducibility Statement

All simulation code, similarity score calculations (IBS, BSI, BSI-VAR), and figure generation were performed using open-source Python libraries. The complete workflow is fully scripted and can be reproduced with standard Python 3.10 environments using the libraries specified above. All random processes (e.g., allele assignment, genotype simulation) were initialized with fixed random seeds where appropriate to ensure reproducibility of results across runs. Custom Python scripts developed for this study are available upon reasonable request to the corresponding authors.

### 2.3 Simulation Design

#### 2.3.1 Random Genotype Simulation for IBS vs Normalized BSI

We simulated genotype data for 100 individuals across 500 bi-allelic SNPs.

Alleles were randomly assigned (‘A’ or ‘G’) based on fixed possibilities (‘AA’, ‘AG’, ‘GG’) without respecting allele frequency distributions. We computed:

IBS: simple genotype match proportion,

Normalized BSI: frequency-weighted matching score using assigned minor allele frequencies (MAFs between 0.01 and 0.5).

Also, we performed 1000 random pairwise comparisons and plotted IBS vs Normalized BSI scores, colored by the number of rare SNP matches (defined as MAF ≤ 0.05).

This first simulation illustrates how frequency weighting prioritizes rare variant matches beyond simple IBS matching.

#### 2.3.2 Forced Rare SNP Matching Simulation for Parent-Child, Siblings, and Random Pairs

To evaluate the performance of IBS and BSI in differentiating related vs unrelated individuals:

We simulated 500 SNPs, assigning random allele frequencies between 0.01 and 0.5. 50 rare SNPs (MAF ≤ 0.05) were force-matched in Parent-Child and Sibling pairs to enrich rare variant sharing.

Three groups were simulated:

##### Parent-Child pairs

Child inherited one allele from Parent at each locus, with forced matching on rare SNPs.

##### Siblings

Independent random inheritance but with rare SNPs force-matched.

##### Random pairs

Genotypes generated completely independently based on MAFs.

Pairwise IBS and BSI scores were computed.

Boxplots were generated comparing the similarity scores across groups.

Mann-Whitney U tests were performed to assess statistical differences between groups.

#### 2.3.3 Large-Scale PCA Simulation with Frequency-Aware Genotype Modeling

To assess discrimination power at scale, 10,000 SNPs were simulated. 500 rare SNPs were injected (MAF ≤ 0.05).

Genotypes for 90 individuals were generated according to SNP-specific allele frequencies:

Parent-Child pairs: simulated by (Mendelian inheritance + rare SNP matching).

Siblings: independent inheritance + rare SNP matching.

Random individuals: generated independently respecting MAFs.

Similarity matrices for IBS, BSI, and BSI-VAR were computed across all individuals. Matrices were mean-centered and PCA was applied to project individuals along PC1 and PC2.

## Results

### 3.1 Random Genotype Simulation (IBS vs Normalized BSI Scatter Plot)

In the first simulation, 100 individuals were randomly genotyped across 500 SNPs without modeling allele frequencies. 1000 random pairwise comparisons were performed, calculating both IBS and Normalized BSI scores. The scatterplot of IBS vs BSI showed a strong correlation overall. However, data points with higher numbers of rare SNP matches (colored yellow and green) tended to deviate above the diagonal IBS = BSI line. This indicates that BSI gives higher similarity scores when rare variants are matched, while IBS remains insensitive. Thus, Normalized BSI successfully prioritizes rare variant matches, a critical property for precision genetic analysis.

**Figure 1:**
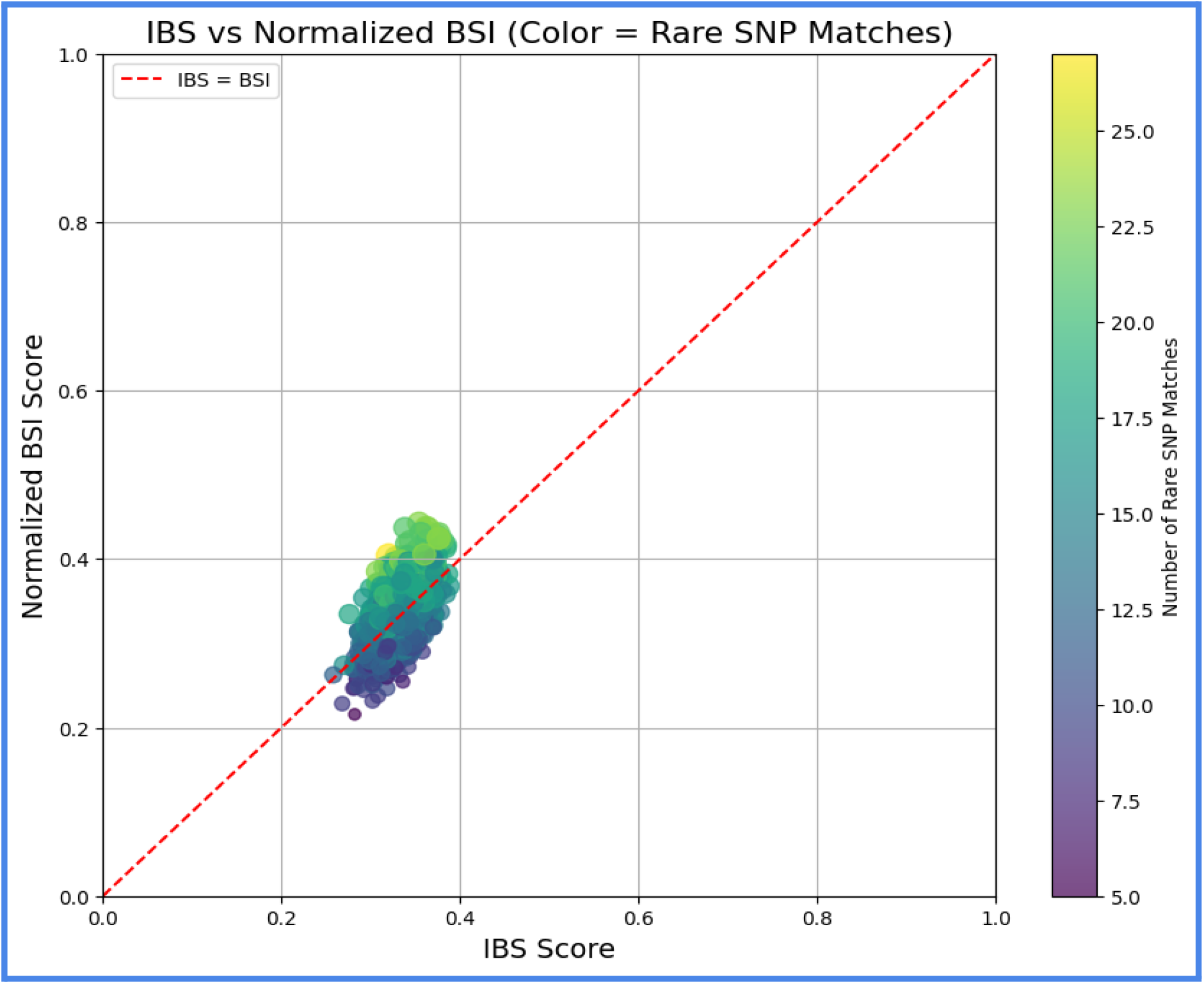
IBS vs BSI Scatter Plot Highlighting Rare Variant Matches

### 3.2 Forced Rare SNP Matching Simulation (Boxplots and Statistical Tests)

In the second simulation, 10 Parent-Child pairs, 10 Sibling pairs, and 10 Random pairs were generated with forced rare SNP matching under controlled allele frequencies.

**Figure 2.**
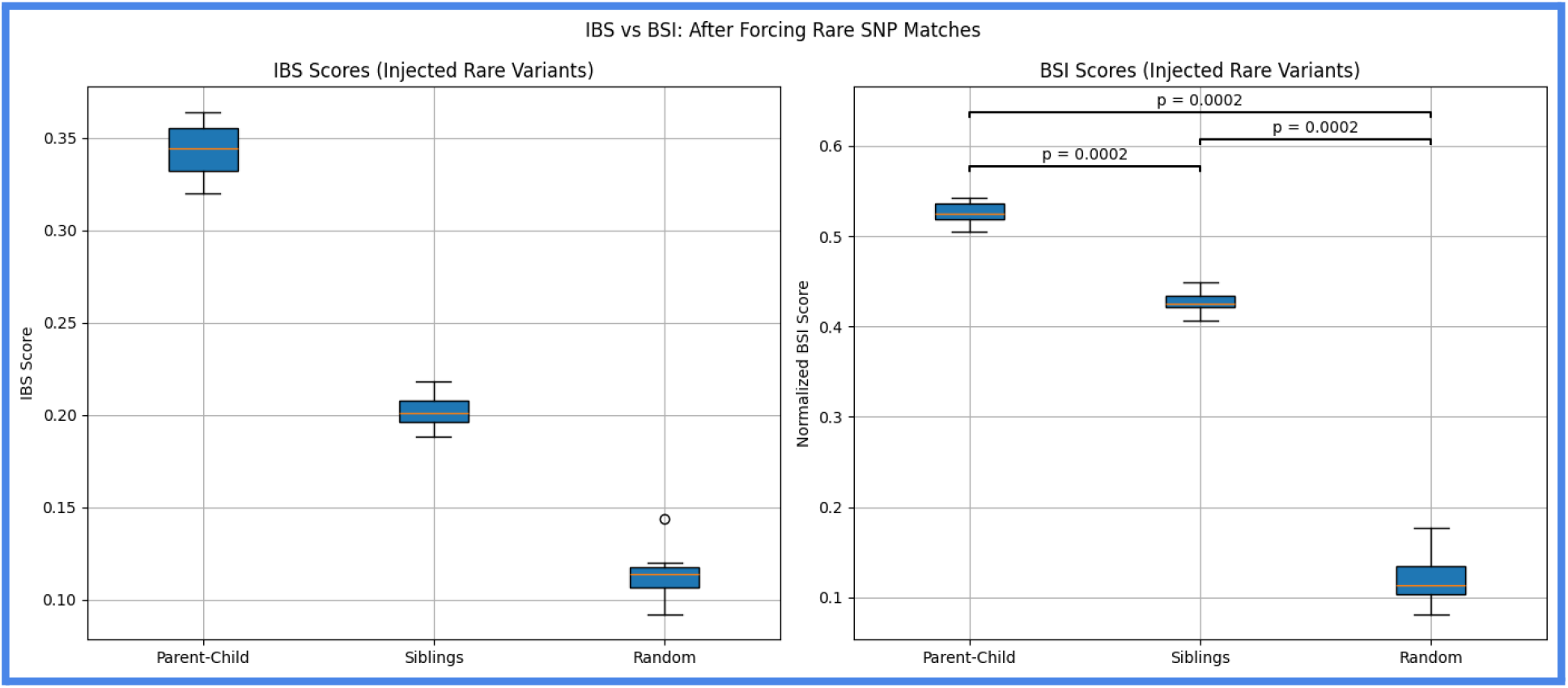
Boxplots Comparing IBS and BSI Scores After Rare Variant Injection. BSI scores provide greater separation between Parent-Child, Sibling, and Random groups compared to IBS, confirmed by lower p-values

#### IBS Summary Stats

Parent-Child : median = 0.3440, mean = 0.3428

Siblings : median = 0.2010, mean = 0.2018

Random : median = 0.1140, mean = 0.1130

#### BSI Summary Stats

Parent-Child : median = 0.5250, mean = 0.5264

Siblings : median = 0.4256, mean = 0.4266

Random : median = 0.1131, mean = 0.1199

Boxplots demonstrated that BSI scores provided greater separation between related and unrelated individuals compared to IBS. Statistical testing confirmed this improvement:

**Table 1.** Summary of Median IBS and Normalized BSI Scores Across Groups.Parent-Child and Sibling pairs show higher BSI scores than Random pairs, highlighting improved discrimination of genetic relationships.

|  |
| --- |
| Parent-Child IBS vs BSI p-value = 0.0002 |
| Sibling IBS vs BSI p-value = 0.0002 |
| Random IBS vs BSI p-value = 0.7336 |

BSI significantly enhanced the ability to discriminate true genetic relationships compared to IBS, while maintaining similar separation among unrelated individuals

### 3.3 Large-Scale PCA Simulation (10,000 SNPs)

In the full simulation of **<u>10,000</u>** SNPs:

IBS PCA: Partial separation, but overlap between Siblings and Random pairs remained.

BSI PCA: Improved separation, with Random individuals moving farther apart from Parent-Child and Sibling groups.

BSI-VAR PCA: Achieved maximum separation, creating three distinct clusters with minimal overlap.

The distance along PC1 between Random and Parent-Child groups was substantially greater for BSI and especially BSI-VAR compared to IBS.

Thus, frequency-aware methods (BSI and BSI-VAR) outperform IBS in both small-scale and large-scale simulations for distinguishing genetic relationships

**Figure 3.**
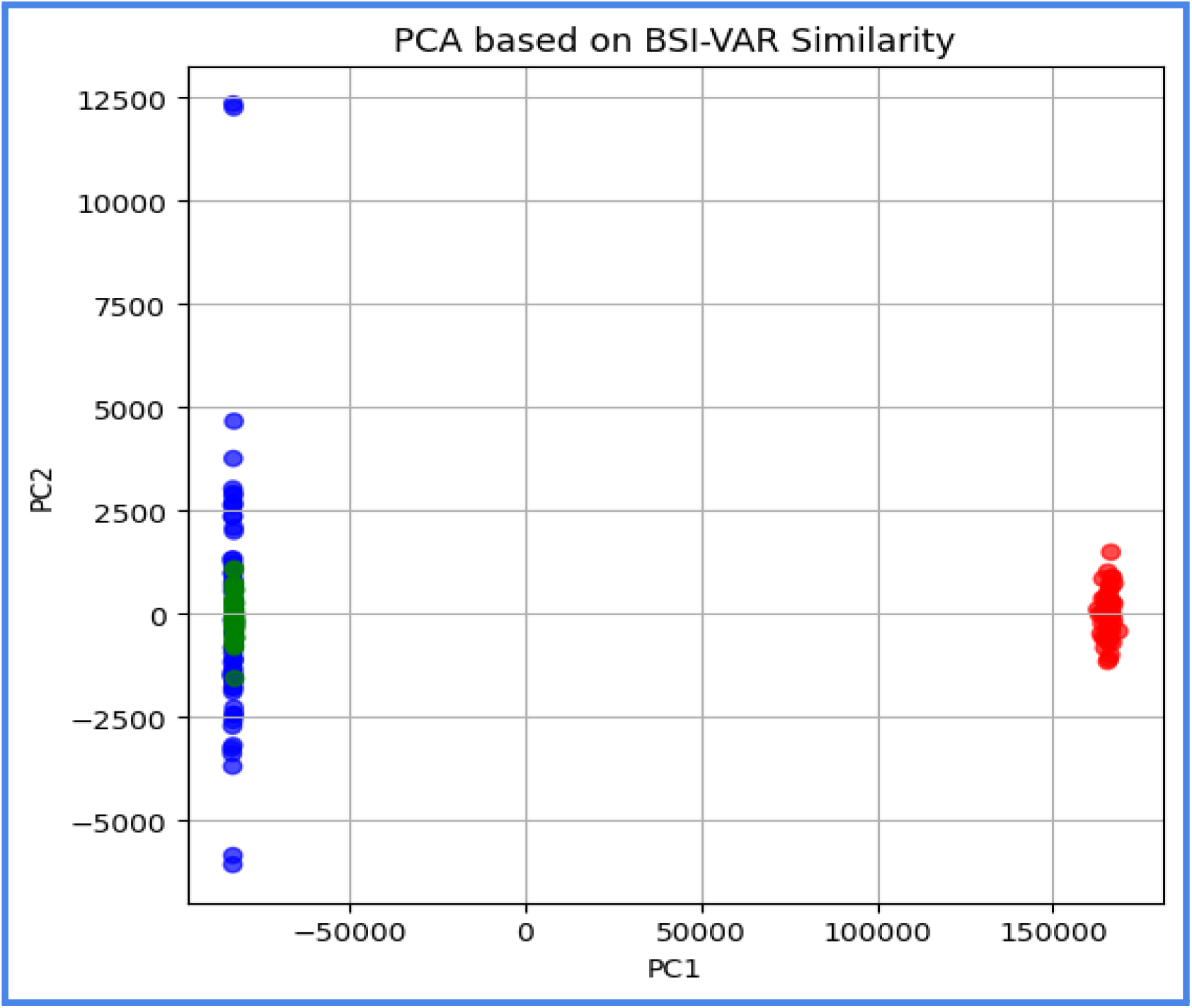
PCA Based on BSI-VAR Similarity Matrix. BSI-VAR achieves the clearest clustering of related pairs, amplifying the contribution of rare variants to genetic similarity assessment.

**Figure 4.**
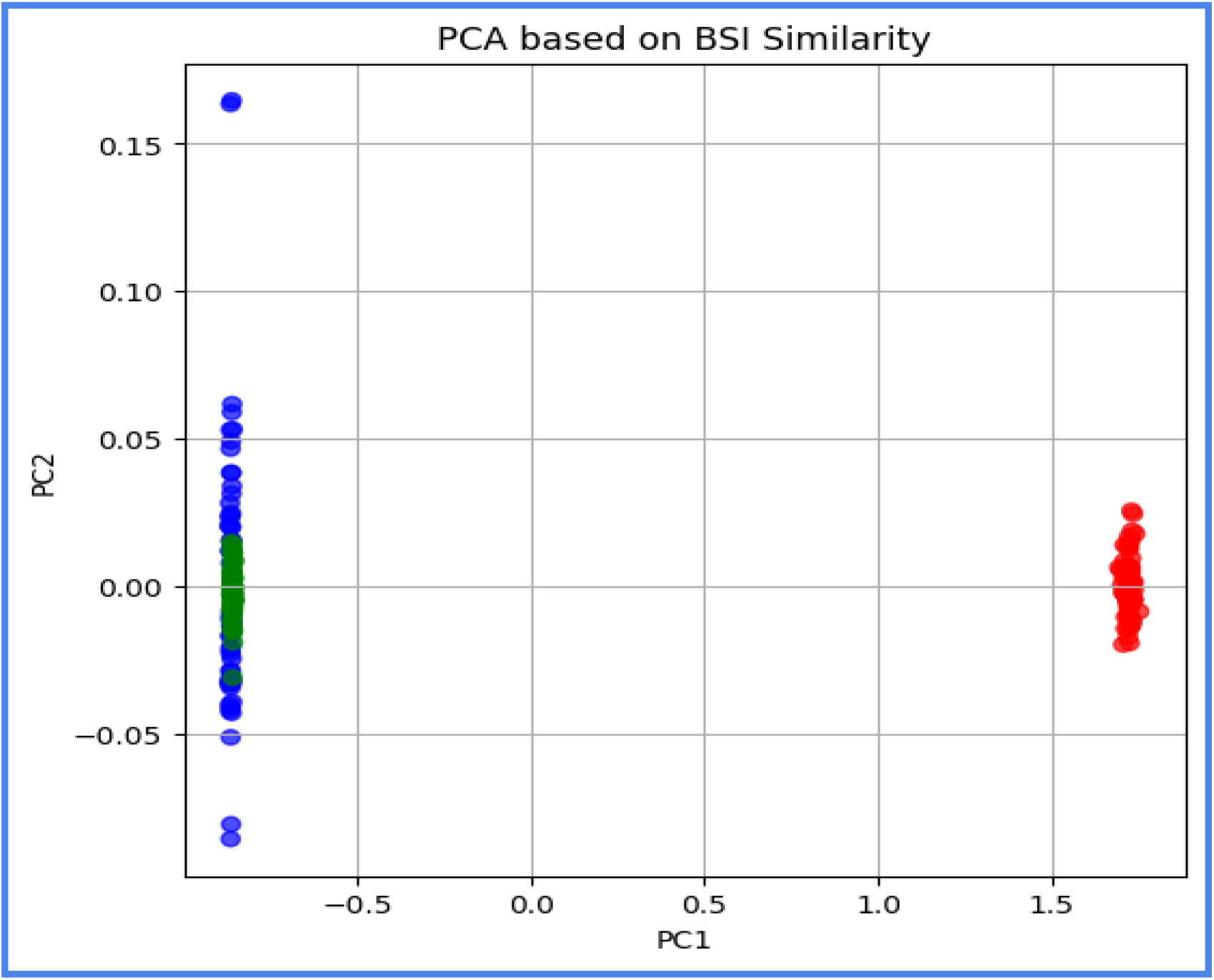
PCA Based on BSI Similarity Matrix. PCA of BSI scores reveals improved separation of Parent-Child and Sibling pairs from Random individuals compared to IBS.

**Figure 5.**
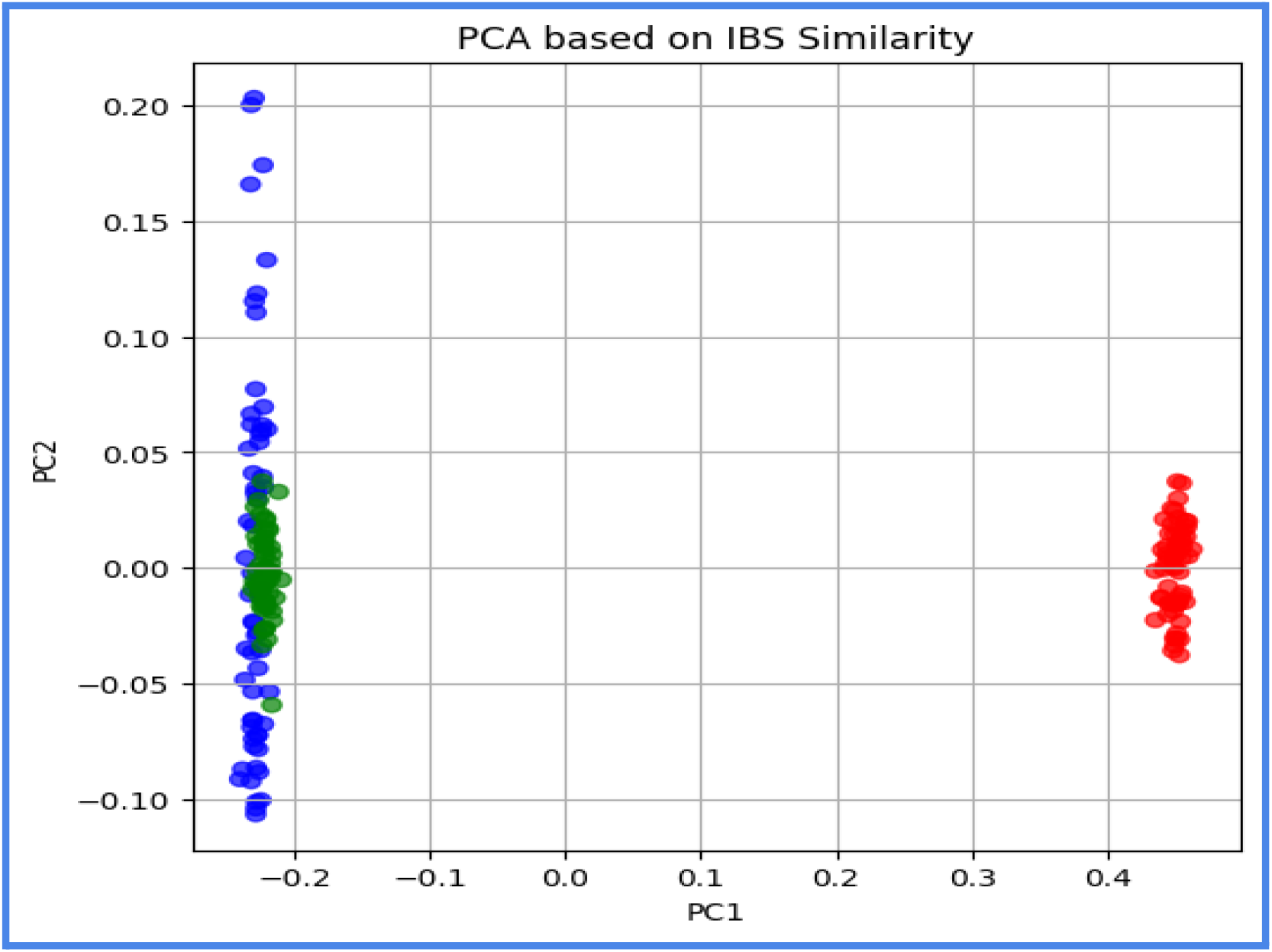
PCA Based on IBS Similarity Matrix. Principal Component Analysis of IBS scores shows partial clustering of related pairs but substantial overlap with random pairs.

## Discussion

Across all simulation scenarios, BSI and BSI-VAR consistently demonstrated superior performance over IBS. In random genotype comparisons, BSI more effectively prioritized rare variant matches, an aspect to which IBS is largely insensitive. In simulations with forced rare variant sharing, BSI provided significantly improved separation between related and unrelated individuals, with consistently lower p-values. In **large-scale** simulations, BSI-VAR achieved the clearest and most robust separation of Parent-Child, Sibling, and Random groups in PCA projections.

These findings highlight that incorporating allele frequency information and accounting for partial genotype sharing meaningfully improves the resolution of genetic relationships. Such improvements are particularly relevant for applications in precision medicine, rare disease gene discovery, and population genetics. By directly addressing the limitations of IBS, BSI and BSI-VAR offer practical, interpretable, and theoretically grounded advances in the field of genomic similarity analysis.

## Conclusion

BSI and BSI-VAR represent a fundamental advance over traditional IBS by incorporating rare variant prioritization and dosage-sensitive weighting, achieving superior resolution of genetic relationships across diverse simulations.

## Novelty Statement

To our knowledge, no prior work has introduced a genetic similarity metric that simultaneously applies:

^1^ Inverse binomial variance weighting based on minor allele frequencies.

^2^ Proportional penalization of allele dosage mismatches.

The Bouziane Similarity Index with Variance Adjustment (BSI-VAR) thus offers a novel, frequency-aware, and dosage-sensitive approach to quantifying genetic closeness, specifically designed to amplify the contribution of rare, stable variants while continuously penalizing partial or complete mismatches.

This advancement addresses critical limitations of classical metrics such as IBS and kinship coefficients, particularly in contexts where rare variant prioritization is crucial, such as precision medicine and population-diverse genomic analyses.

## Future Directions

Future directions include extending the Bouziane Similarity Index (BSI) framework to accommodate multi-allelic variants, structural variations, and imputed genotype data. Empirical validation using large-scale biobank and sequencing datasets will be essential to evaluate performance across diverse ancestries and technical platforms.

Further, integrating frequency-aware similarity scores into machine learning models may unlock new opportunities for rare variant discovery, relationship inference, and personalized medicine.

## Acknowledgment

The authors sincerely thank Dr. CHOAYB DJEDDI for his valuable support and encouragement throughout this work.

The authors also acknowledge genomeDZ for supporting the computational work associated with this study.

## Notes

### Competing Interest Statement

The authors have declared no competing interest.

